# High Maternal Adiposity During Pregnancy Programs an Imbalance in the Lipidome and Predisposes to Diet-induced Hepatosteatosis in the Offspring

**DOI:** 10.1101/2023.03.06.531438

**Authors:** Taylor B. Scheidl, Jessica L. Wager, Larissa G. Baker, Amy L. Brightwell, Katrina M. Melan, Sebastian Larion, Ousseynou Sarr, Timothy RH. Regnault, Stefan J Urbanski, Jennifer A. Thompson

## Abstract

**Background:** Exposure to high maternal adiposity *in utero* is a significant risk factor for the later-life development of metabolic syndrome (MetS), including non-alcoholic fatty liver disease (NAFLD). We have previously shown that high pre-pregnancy adiposity programs adipose tissue dysfunction in the offspring, leading to spillover of fatty acids into the circulation, a key pathogenic event in obesity-associated MetS. Herein, we hypothesized that programming of adipose tissue dysfunction in offspring born to overweight dams increases the risk for developing NAFLD.

**Results:** Females heterozygous for leptin receptor deficiency (Het_*db*_) were used as a model of high pre-pregnancy adiposity. Wild-type (Wt) offspring born to Het_*db*_ pregnancies gained significantly more body fat following high fat/fructose diet (HFFD) compared to Wt offspring born to Wt dams. HFFD increased circulating free fatty acids (FFA) in male offspring of control dams, while FFA levels were similar in HFFD-fed offspring from Wt dams compared to CD or HFFD-Wt offspring from Het_*db*_ dams. Despite female-specific protection from diet-induced FFA spillover, both male and female offspring from Het_*db*_. dams were more susceptible to diet-induced hepatosteatosis. Lipidomic analysis revealed that CD-offspring of overweight dams had decreased hepatic PUFA levels compared to control offspring. Changes to saturated fatty acids (SFA) and the *de novo* lipogenic (DNL) index were diet driven; however, there was a significant effect of the intrauterine environment on FA elongation and Δ9 desaturase activity.

**Conclusion:** High maternal adiposity during pregnancy programs a susceptibility to diet-induced hepatosteatosis.

## INTRODUCTION

Non-alcoholic fatty liver disease (NAFLD) is the hepatic manifestation of the metabolic syndrome (MetS) and has grown in prevalence from ~25% to nearly 40% over the past couple of decades, primarily due to rising obesity rates (1). Consequent to the declining age of obesity-onset, a greater number of women are obese at reproductive age, approximately 25% of women in the USA (2). Human studies have established a relationship between maternal obesity and later life development of components of MetS in the offspring, including obesity, insulin resistance and dyslipidemia (3,4). A recent study in a Swedish cohort of young adults revealed that being born to an obese mother increased the risk for developing NAFLD in adulthood by 3-fold (3). Therefore, exposure to an abnormal intrauterine milieu may be an important, but underappreciated risk factor for NAFLD.

Adipose tissue function distinguishes metabolically healthy obesity from unhealthy obesity. Dysfunctional adipose tissue is characterized by chronic inflammation, insulin resistance, dysregulated lipolysis, and a failure to buffer excess lipids, leading to lipid spillover and ectopic deposition in peripheral organs such as the liver. Therefore, adiposopathy is a key trigger for the onset of obesity-associated lipid accumulation in the liver. Recently, we showed that high prepregnancy adiposity programs an increased risk for diet-induced adiposopathy in the offspring, in a sex-specific manner (5). While an effect of the intrauterine environment on diet-induced perturbations in adipose tissue function and lipid homeostasis was observed in both male and female offspring born to overweight dams, females from normal pregnancies were protected from the cardiometabolic effects of high fat diet. In the current study, we sought to determine the influence of the intrauterine environment and sex on diet-induced hepatic steatosis and lipid metabolism.

## METHODS

### Animals

All experimental protocols were approved by the University of Calgary Animal Care Committee (AC17-0149) and conducted in accordance with guidelines by the Canadian Council on Animal Care Ethics. Female mice heterozygous for deletion of the leptin receptor (Het_*db*_) (Jackson Laboratory, stock #: 000697) were used as a model of high pre-pregnancy adiposity, as characterized in our previous study. Virgin 14-week old Het_*db*_ or wild-type (Wt) C57BL/6J females were mated with Wt males. Following delivery, offspring of Het_*db*_ dams were genotyped by qPCR and only Wt offspring used to isolate the effect of the *in-utero* environment. Pups were weaned on post-natal day (PND) 21. At 7-weeks of age, offspring were randomly assigned to a high-fat/high-fructose (HFFD) (Research Diets Inc. D08040105I) or control diet (CD) (Research Diets Inc. D12450Ki) for 15 weeks. At 22-30 weeks of age, mice were induced under isoflurane inhalation and euthanized by decapitation. Blood was collected in heparinized syringes and plasma collected by centrifugation. Livers were collected and either flash frozen in liquid nitrogen or fixed in 4% paraformaldehyde for sectioning.

### Body composition and fasting NEFA

Prior to euthanasia, body composition was measured by time domain nuclear magnetic resonance (TD-NMR) spectroscopy (LF90II, Bruker). Whole-body fat and lean body mass were expressed as a % of body weight. Fasting NEFA levels were measured using a colorimetric assay (Abcam) according to the manufacturer’s instructions. Samples were run in triplicate and absorbance read on a SpectraMax ID3 microplate reader.

### Histopathology

Fixed hepatic tissue was embedded in paraffin and 4μm sections mounted on slides by the University of Calgary Faculty of Veterinary Medicine Diagnostic Services Unit. Sections were stained with hematoxylin and eosin (H&E) or picrosirius red stain. Steatosis, inflammation, ballooning, and fibrosis were evaluated blind by an experienced histopathologist using the steatosis, activity, fibrosis (SAF) scoring system. Grades 0-3 of steatosis were defined as <5%, 5-33%, 33-66% and >33% steatosis, respectively. Inflammation was scored as 0 (absence of foci), 1 (≤ 2 foci) or 2 (>2 foci). Hepatocyte ballooning was absent (grade 1), apparent by clusters of enlarged hepatocytes with rarified cytoplasm (grade 2) or with more severe hepatocyte enlargement (grade 3). Fibrosis scores 0-3 were defined as the absence of fibrosis, perisinusoidal or periportal fibrosis, perisinusoidal and periportal fibrosis, bridging fibrosis and cirrhosis, respectively. Representative images of liver sections were captured on a Nikon TI Eclipse brightfield microscope.

### RNA extraction and qPCR

Total RNA was isolated from flash frozen liver using the RNeasy Mini Kit, assessed for integrity using a TapeStation assay (University of Calgary Genomic Services), and quantified by nanophotometer (Implen). After digestion of DNA, cDNA was generated from 2μg RNA using the High-Capacity Reverse Transcription kit (Applied Biosystems). Samples were assayed in triplicate for RT-qPCR using SYBR Green master mix (A25741, Applied Biosystems). Amplification was performed using the following settings: 50°C x 2 minutes, 95°C x 2 minutes, 95°C x 1 second, and 58.9°C x 30 seconds for 40 cycles on the QuantStudio 5 Real Time PCR System. Primers were designed using NCBI and validated with melt curve analysis (Table 1). Albumin was used as a reference gene for calculation of mRNA expression.

**Table 1.**
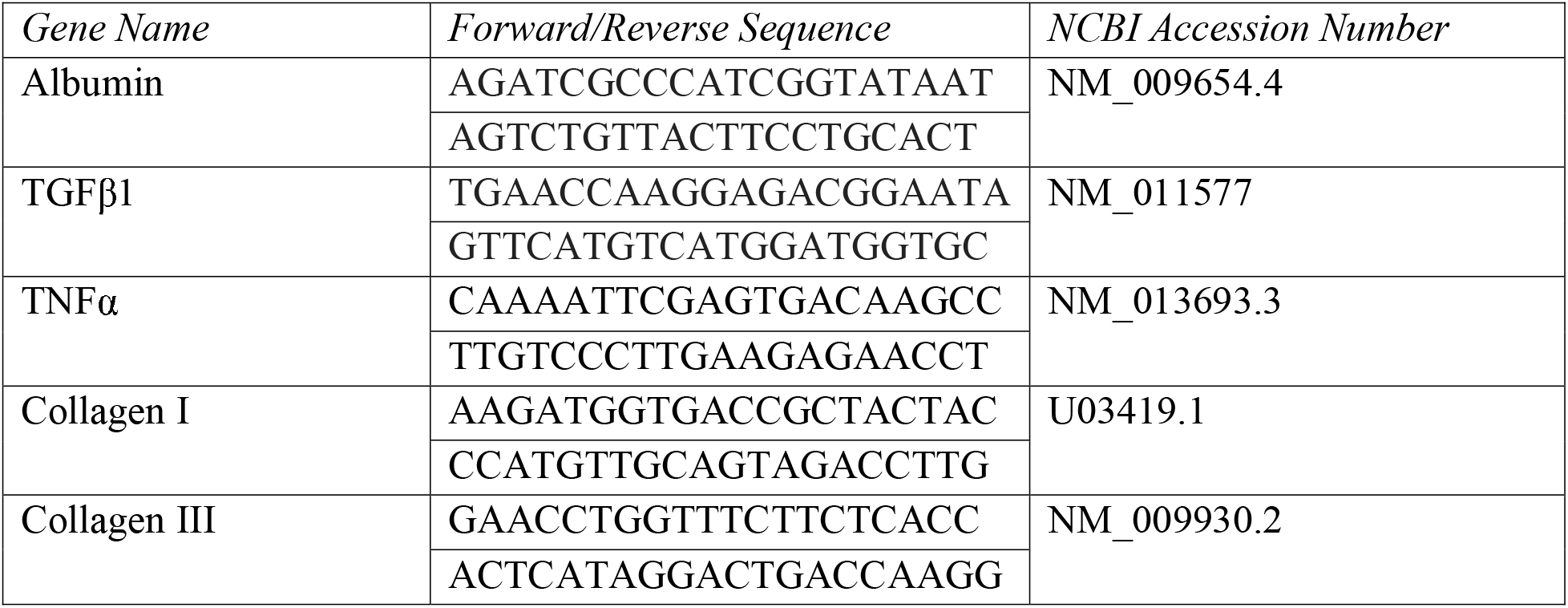
Primer Sequences.

### Lipid extraction and thin layer chromatography for quantification of fatty acid species

Lipid extraction and thin layer chromatography was performed by the Vanderbilt University Medical Center Lipid Lab. Briefly, thin layer chromatography was used to separate lipids extracted using the Folch method. Individual lipid classes were visualized by rhodamine 6G and methylated for extraction. Methylated fatty acids were identified by comparing the retention times to known standards. Odd chain fatty acid standards were used for lipid quantification.

### Indirect measurement of lipogenic enzyme activity

The Δ9 desaturation index was calculated as the ratio of precursors [palmitic acid (C16:0), stearic acid (C18:0)] to products [palmitoleic (C16:1), oleic acid (C18:1)] and reflects the activity of stearoyl-CoA 9-desaturase-1 (SCD1), an enzyme that catalyzes the rate limiting step in the biosynthesis of substrates (C16:1 & C18:1) for incorporation into triglycerides and cholesterol. The *de novo* lipogenesis (DNL) index, reflecting the rate of endogenous FA biosynthesis, was calculated as the ratio of palmitic acid (C16:0) to linoleic acid (C18:2n6). The ratio of C18:0-to-C16:0 was calculated to reflect the elongation index, reflecting the chain elongation of C16:0 catalyzed by fatty acid elongase 6 (Elov6). The sums of MUFA, SFA and PUFA were calculated, and levels expressed as a % of total lipid species.

### Statistical Analysis

Litter effects were controlled by including 1-2 male and female mice per litter in each analysis. Two-way ANOVA was used to determine main effects of diet and intrauterine environment (offspring group), while Sidak’s multiple comparison test in *post-hoc* analysis determined differences between groups. A contingency table was created with histological scoring counts for each group and differences between groups determined using the Chi-squared test. Analyses were performed using GraphPad version 9.0. Comparisons were considered significant when p<0.05 and data are expressed as mean ± SEM.

## RESULTS

### High pre-pregnancy adiposity programs an obesogenic phenotype in the offspring

After 15-weeks of HFFD, whole-body fat mass was similarly increased in males and females (Figure 1. A&B). Fat mass was higher (Figure 1 A&B) and lean body mass lower (Figure 1 C&D) in both female and male HFFD-fed offspring from Het_*db*_ dams vs. Wt dams, and in females there was a significant effect of intrauterine environment on fat mass. In males, HFFD lead to an increase in fasting FFA, while plasma FFA were similar in HFFD-fed male offspring born to Wt dams and Het_*db*_ offspring fed a CD or HFFD (Figure 1E). There was no impact of diet or intrauterine environment on plasma FFA levels in females (Figure 1F).

**Figure 1:**
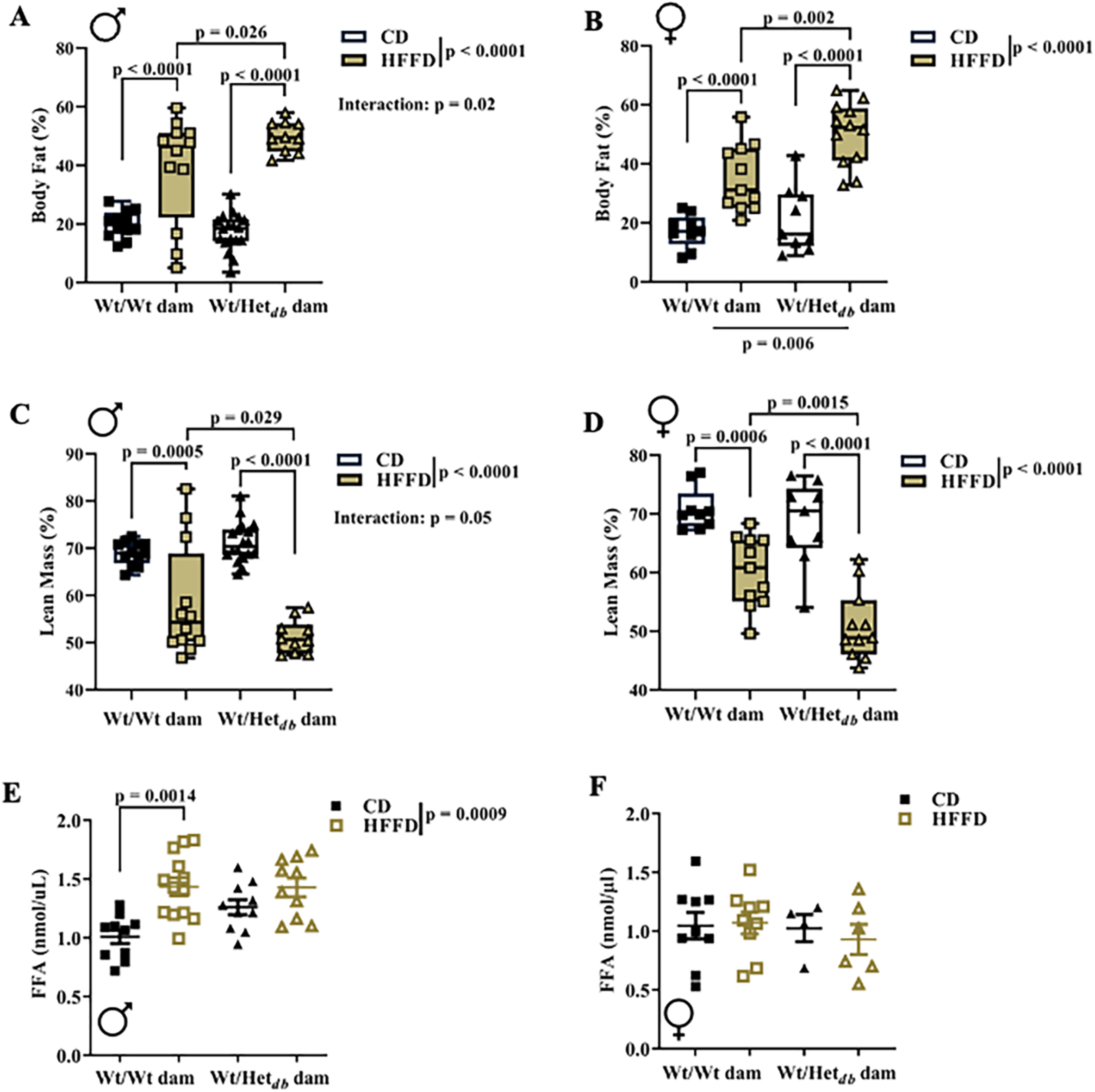
High pre-pregnancy adiposity programs an obesogenic phenotype in the offspring. Body composition was measured by TD-NMR in 5-month old Wt offspring born to Wt (Wt/Wt dam) or Het_*db*_ (Wt/Het_*db*_) dams after 15 weeks of control diet (CD) or high fat/fructose diet (HFFD). Fat mass in A) male and B) female offspring. Lean body mass in C) male and D) female offspring. Fasting levels of plasma free fatty acids were measured in E) male and F) female offspring. Main effects of intrauterine environment and diet were determined by two-way ANOVA, with Sidak’s multiple comparison test to determine differences between groups.

### Offspring born to overweight dams are more susceptible to diet-induced hepatic steatosis

In 5-month-old CD-fed offspring born to Wt or Het_*db*_ dams, no steatosis or hepatocellular ballooning was observed in males. Grade 3 steatosis was observed in 43% and 11% of samples in CD-fed females born to Wt and Het_*db*_ dams, respectively, while grade 2 was observed in 11%-14% of female CD samples. Grade 1 and 2 steatosis was observed in 14% and 29%, respectively, in HFFD-fed control males; while in HFFD-fed control females, 33% of samples were grade 2 and 50% were grade 3. In Chi-square analyses, HFFD had a significant effect on hepatic steatosis in only offspring born to Het_*db*_ pregnancies, and this was the case for both sexes (Figure 2 A-C). Hepatocellular ballooning was observed in only HFFD-fed males and HFFD-fed female offspring from Het_*db*_ dams, although there were no significant effects of diet or intrauterine environment (Figure 2 D&E). There was no apparent fibrosis in any male groups, whereas grade 1 or 2 fibrosis was noted in a few female samples with no significant differences between groups. Similarly, no differences in scoring for inflammation were noted. In male offspring, there was a significant effect of diet on the expression of transforming growth factor beta (*Tgfβ1*), with a significant difference between Wt males born to Het_*db*_ dams on a CD vs. HFFD (Figure 3E). In female offspring, there was a significant effect of diet on hepatic expression of pro-collagen I (*PcI*) and pro-collagen III (*PcIII*), with a significant difference between Wt females born to Het_*db*_ dams on a CD vs. HFFD (Figure 3 B). The expression of tumor necrosing factor alpha (*Tnfα*) was not affected by diet or intrauterine environment.

**Figure 2:**
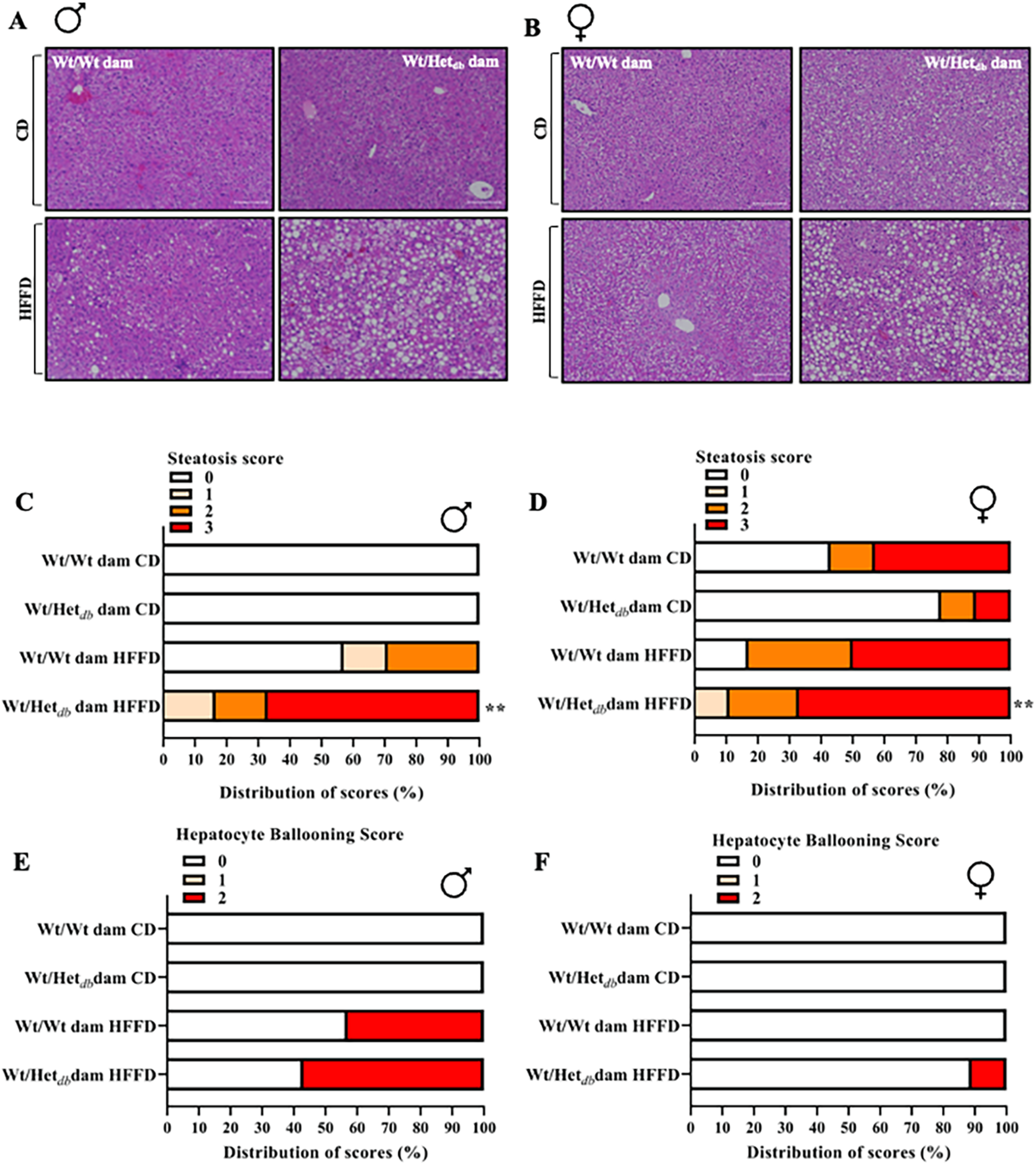
Offspring born to overweight dams are more susceptible to diet-induced hepatic steatosis. After 15 weeks of control diet (CD) or high fat/fructose diet (HFFD), H&E-stained liver sections from Wt offspring born to Wt (Wt/Wt dam) or Het_*db*_ (Wt/Het_*db*_) dams were evaluated by an experienced pathologist. Representative H&E-stained sections of the liver from A) male and B) female offspring. The distribution of scores for steatosis (0–3) in C) male and D) female offspring. The distribution of scores for hepatocellular ballooning (0–2) in E) male and F) female offspring. A contingency table was created to perform a chi-square test to determine differences in score count between groups (n = 6-9/group). ** p < 0.01 Wt/Het_*db*_ offspring CD vs. HFFD.

**Figure 3:**
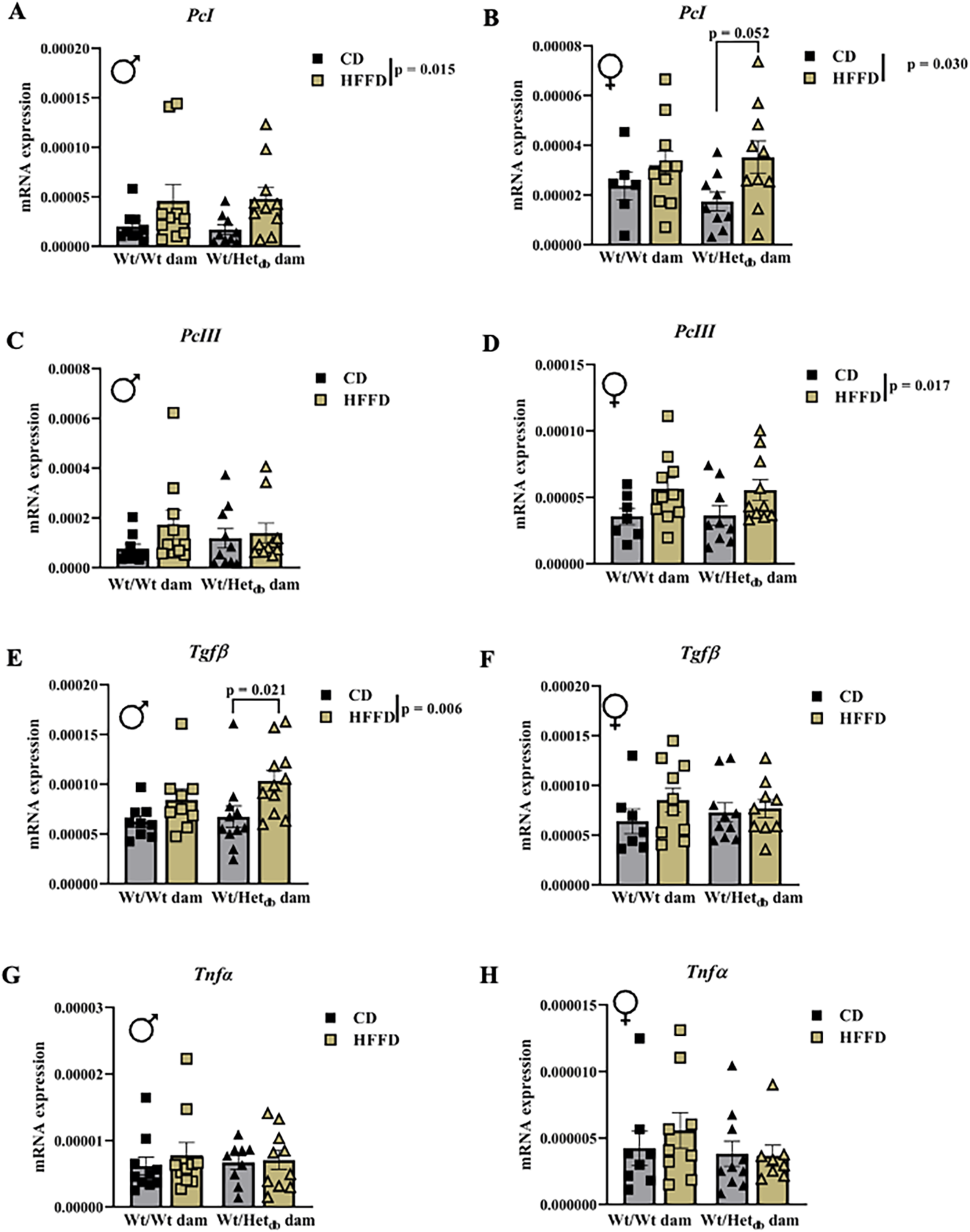
Expression of pro-fibrotic and inflammatory genes in the liver. qPCR was used to measure the expression of A,B) pro-collagen I (*PcI);* C,D) pro-collagen III (*PcIII*), transforming growth factor beta (*Tgfβ);* and tumor necrosing factor alpha (*Tnfα*) in livers of control diet (CD) or high fat/fructose diet (HFFD)-fed 5-month old Wt male and female offspring born to Wt (Wt/Wt dam) or Het_*db*_ (Wt/Het_*db*_) dams. Differences were assessed by two-way ANOVA with intrauterine environment and diet as main effects, while Sidak’s multiple comparison test determined differences between groups.

### Individual SFA species were increased and PUFA decreased by HFFD and in offspring born to overweight dams

Individual lipid species present in abundance in the FFA fraction are shown in Figures 4A & B. Except for 18:1ω9, all major lipid species were affected by the diet [16:0 (p < 0.0001); 18:0 (p < 0.0001); 16:1 (p = 0.0002); 18:1ω7 (p < 0.0001); 18:2 (p < 0.0001); 20:4 (p = 0.023)]. In post hoc analysis, HFFD increased the abundance of 16:0 (p < 0.0001), 18:0 (p < 0.001), 16:1 (p = 0.0007) and 18:1ω7 (p = 0.0005) in Wt offspring born to Wt dams. Similarly, in Wt offspring born to Het_*db*_ dams, there was an increase in 16:0 (p < 0.0001), 18:0 (p < 0.0001), and 18:1ω7 (p < 0.001). In both groups 18:2 was decreased by HFFD (p < 0.0001). There was a main effect of the intrauterine environment on 16:0 (p = 0.0027), 18:1ω7 (p = 0.046), 18:2 (p = 0.0032) and 20:4 (p = 0.018). In CD-fed offspring, 16:0 was increased (p = 0.0015) and 18:2 decreased (p = 0.0008) in Het_*db*_ vs. Wt offspring. The increase in 18:0 after HFFD was greater in offspring from Het_*db*_ vs. Wt pregnancies (p = 0.0271). There was a significant interaction between diet and the intrauterine environment for 16:0 (p = 0.0117) and 18:2 (p = 0.004). For minor lipid species present in low abundance (Figure 4 C & D), 18:3ω6, 18:3ω3, 20:3ω6 and 20:5 were all below the detection limit in both HFFD groups. Comparing CD-fed offspring from Het_*db*_ vs. Wt dams, 18:3ω6 (p = 0.0089), 18:3ω3 (p = 0.0058) and 20:5 (p = 0.046) were all lower in Het_*db*_ offspring.

**Figure 4:**
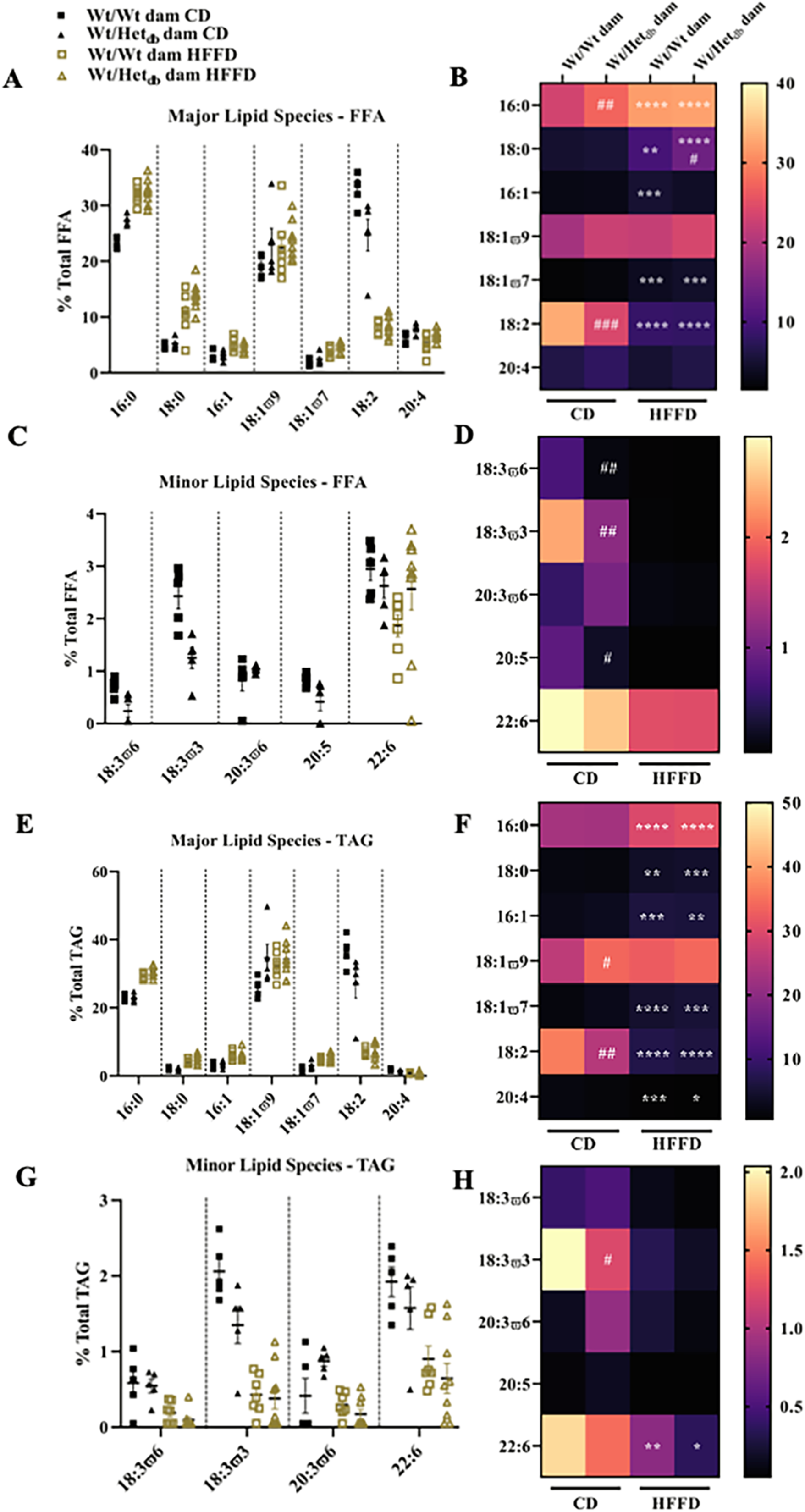
Individual FFA and TG species in hepatic lipid fractions. The Folch method was used to extract lipids from liver samples of 12-week-old male offspring for the separation and identification of individual lipid classes by thin layer chromatography. A) Major lipid species present in abundance in FFA, including saturated FA: palmitic acid (16:0) and stearic acid (18:0); monounsaturated FA: palmitoleic acid (16:1), oleic acid 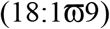 and vaccenic acid 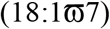; and polyunsaturated FA: linoleic acid (18:2) and arachidonic acid (20:4). B) Heat map showing geometric means of major individual FFA lipid species. C) Minor lipid species present in trace amounts in FFA, including polyunsaturated FAs: γ-linolenic acid (18:3ω6), α-linolenic acid (18:3ω3), dihomo-γ-linolenic acid (20:3ω6) and docosahexaenoic acid (22:6). Heat map showing geometric means of minor individual FFA lipid species. E) Major lipid species present in TG. F) Heat map showing geometric means of minor individual TG species. Main effects of intrauterine environment and diet were determined by two-way ANOVA, with Sidak’s multiple comparison test to determine differences between groups (n = 5-9/group). * p < 0.05 HFFD vs. CD (same intrauterine environment); # p < 0.05 Wt/Wt dam vs. Wt/Het_*db*_ dam (same diet).

Major individual lipid species in the TAG fraction are shown in Figures 4E & F. Similar to the FFA fraction, diet had an effect on all major lipid species except for 18:1ω9 [16:0 (p < 0.0001); 18:0 (p > 0.0001); 16:1 (p < 0.0001); 18:1ω7 (p < 0.0001); 18:2 (p < 0.0001); 20:4 (p < 0.0001)]. HFFD increased the abundance of 16:0 (p < 0.0001); 18:0 (p = 0.001), 16:1 (p < 0.0001), and 18:1ω7 (p < 0.0001) in offspring from Wt dams. In Het_db_ offspring, HFFD also increased the presence of 16:0 (p < 0.0001), 18:0 (p < 0.0001), 16:1 (p = 0.0018), and 18:1ω7 (p = 0.0002). The abundance of 18:2 was reduced by HFFD in Wt and Het_db_ offspring (both p < 0.0001). Similarly, 20:4 was decreased with HFFD in both Wt (p = 0.004) and Het_db_ (p = 0.023) offspring. There was an effect of the intrauterine environment on 18:1ω9 (p = 0.028), 18:1ω7 (0.0265) and 18:2 (p = 0.0126). *Post hoc* analysis showed that in CD-fed groups, there was an increase in 18:1ω9 (p = 0.0393) and decrease in 18:2 (p = 0.0056) in Het_db_ vs. Wt offspring. For minor lipid species (Figure 1G & H), >40% of samples were below the detection limit in the HFFD groups for 18:3ω6 and 18:3ω3 and thus comparisons were not made. In CD-fed offspring, there was a decrease in the abundance of 18:3ω3 in Het_*db*_ vs. Wt (p = 0.0433). There was a significant effect of the diet on the abundance of 22:6 (p < 0.0001) with a HFFD-associated decrease in Wt (p = 0.0073) and Het_db_ (p = 0.0103) offspring. No comparisons were made for 20:3ω6 due to too many samples below the detection limit, and 20:5 was undetected in the TAG fraction.

### The activity of lipogenic enzymes is perturbed by HFFD and in offspring born to overweight dams

Lipidomic profiles were used to indirectly measure the activity of lipogenic enzymes. The Δ9 fatty acid desaturase, SCD1, catalyzes the conversion of palmitic acid (16:0) and stearic acid (18:0) to palmitoleic acid (16:1) and and oleic acid (18:1η9), respectively, key substrates used in the biosynthesis of complex lipids. The ratio of products-to-precursors was calculated to reflect the activity of SCD1. There was a significant effect of diet on the Δ9 desaturation index, with 16:1/16:0 increased in HFFD vs. CD control offspring, and a decrease in the 18:1/18:0 due to HFFD in both groups of offspring (Figure 5A&B). The 16:0/16:1 ratio was not different between control offspring fed a HFFD and offspring born to Het_*db*_ pregnancies fed either a CD or HFFD. The 18:1/18:0 ratio was higher in CD-fed offspring from Het_*db*_ dams vs. Wt dams and there was a significant effect of intrauterine environment on 18:1/18:0. The rate of DNL was inferred from the abundance of 16:0, the main product of DNL, relative to the essential FA, linoleic acid (18:2). There was a significant effect of diet on the DNL index, with increases in HFFD vs. CD offspring from both groups (Figure 5C). Elongation of 16:0 to 18:0, catalyzed by Elov6, was significantly affected by diet, and increased in HFFD vs. CD in both groups of offspring (Figure 5E). The elongation index was increased in CD-fed offspring from Het_*db*_ dams vs. Wt dams.

**Figure 5:**
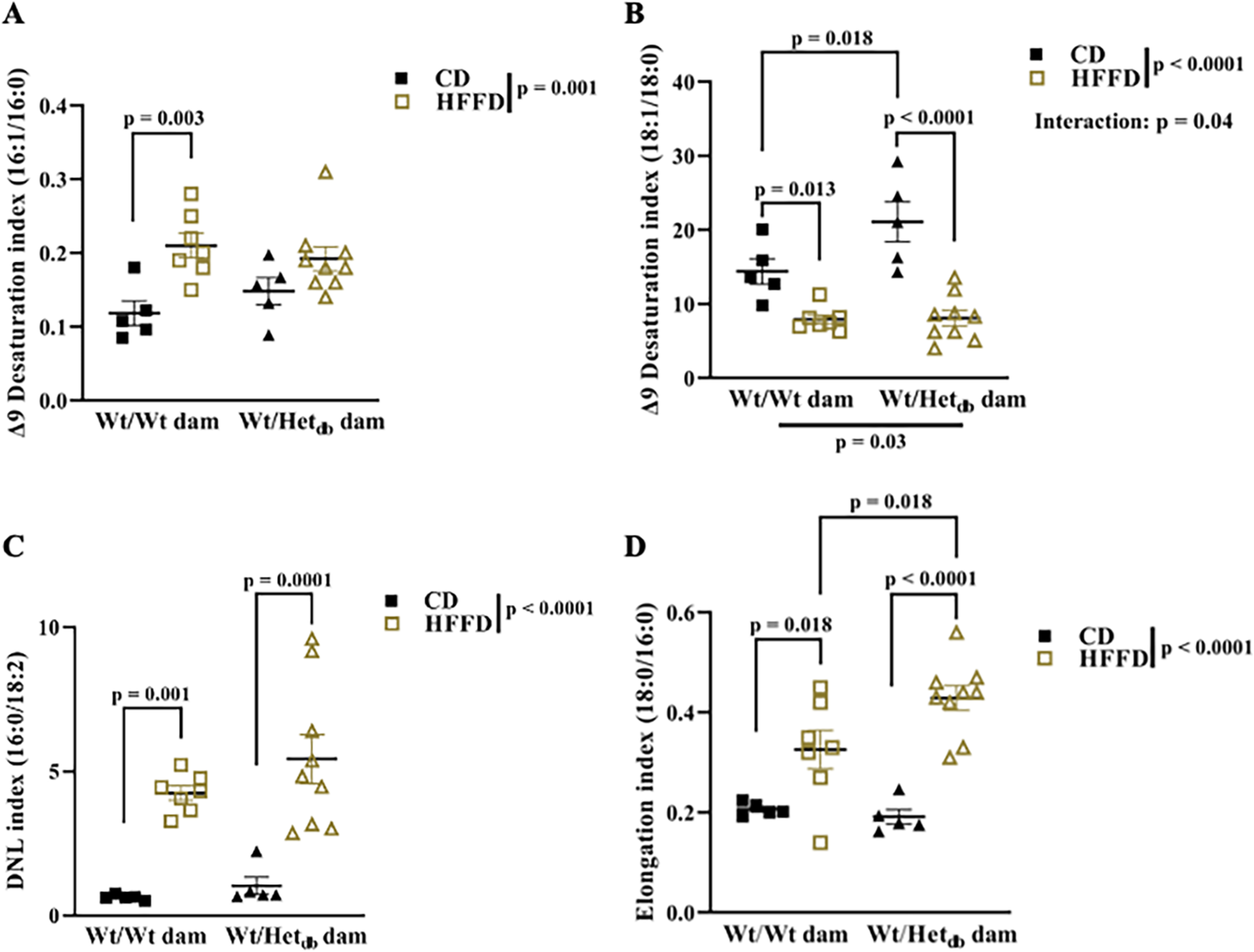
Effects of intrauterine environment and diet on lipogenic enzyme activity. Lipidomic profiles of livers collected from 12-week-old male offspring fed a control diet (CD) or high fat/fructose diet (HFFD) were used to indirectly measure the activity of lipogenic enzymes. The Δ9 desaturation index, reflecting stearoyl-CoA desaturase activity, was calculated as the ratios of A) palmitoleic acid (C16:1) to and palmitic acid (16:0) and B) oleic acid (18:1) to stearic acid (18:0). C) The *de novo* lipogenic (DNL) index was calculated as the ratio of palmitic acid (16:0) to linoleic acid (18:2). D) The elongation index was calculated as the ratio of stearic acid (18:0) to palmitic acid (16:0). Main effects of intrauterine environment and diet were determined with two-way ANOVA with Sidak’s multiple comparison test to determine differences between groups (n = 5-9).

### HFFD and an abnormal intrauterine milieu disturbs the balance in hepatic MUFA and PUFA

The total FFA abundance of MUFA, SFA and PUFA in FFA were significantly affected by diet. SFA were increased and PUFA decreased by HFFD in both groups of offspring (Figure 6B&C). Total MUFA were increased in HFFD vs. CD offspring from control pregnancy, while there were no differences in total MUFA between HFFD-fed control offspring and HFFD or CD-fed offspring from Het_*db*_ pregnancies (Figure 6A). Total PUFA were lower in CD-fed offspring from Het_*db*_ dams compared to CD-fed offspring from Wt dams (Figure 6C).

**Figure 6:**
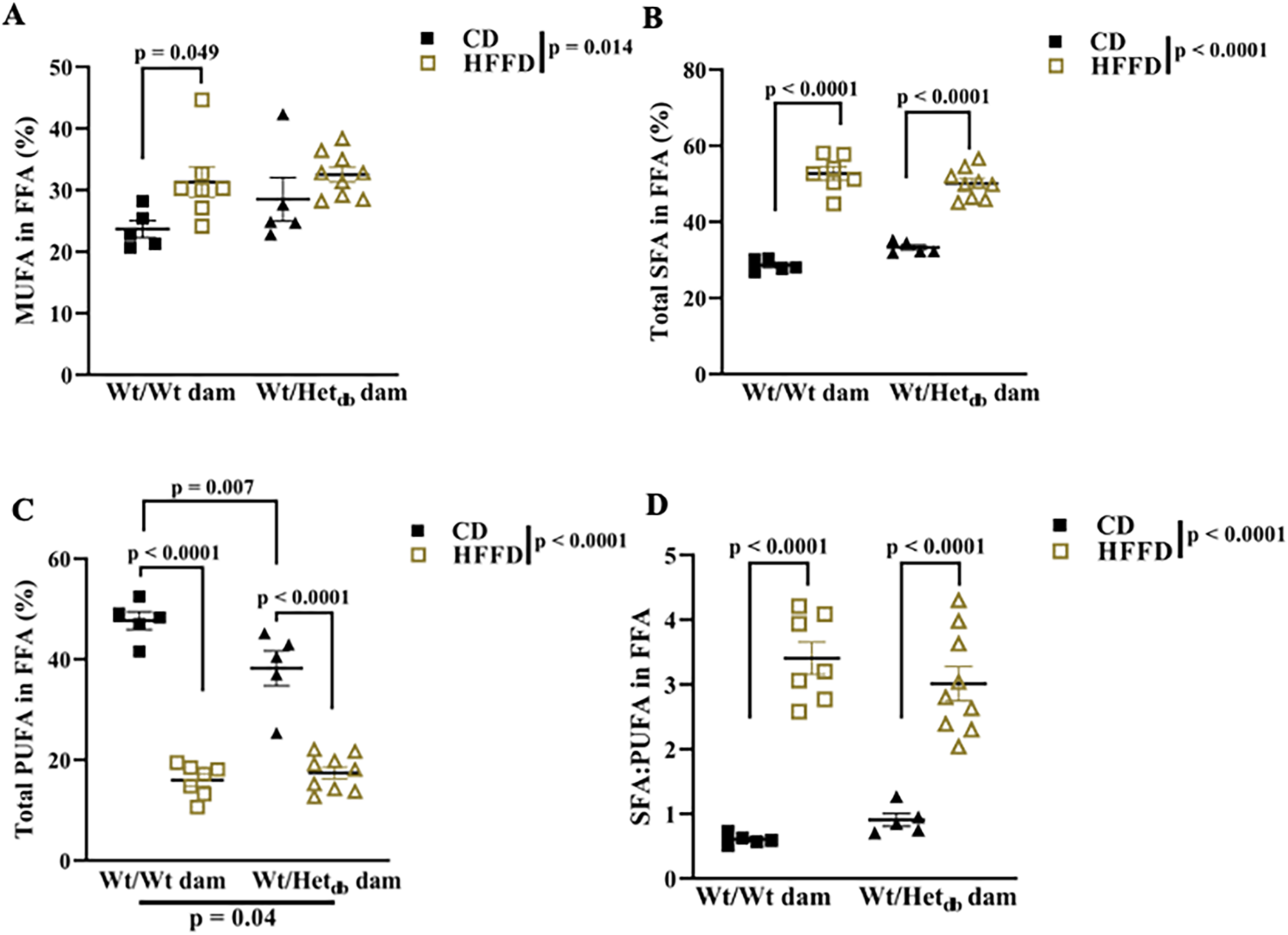
Effects of intrauterine environment and diet on MUFA, SFA and PUFA in hepatic lipid fractions. The sum of A) monounsaturated fatty acids (MUFA), B) saturated fatty acids (SFA) and C) polyunsaturated fatty acids (PUFA) were calculated and expressed as a % of total lipid species. D) The ratio of the sum of SFA relative to the sum of PUFA. Main effects of intrauterine environment and diet were determined by two-way ANOVA with Sidak’s multiple comparison test to determine differences between groups (n = 5-9).

## DISCUSSION

It is well established that maternal health during pregnancy shapes lifelong risk for chronic disease in the offspring. Several human studies have demonstrated that babies born to overweight or obese mothers are more likely to develop NAFLD and other components of the metabolic syndrome later in life (3,6). Herein, we used female mice heterozygous for leptin receptor deficiency as a model of maternal metabolic dysfunction. Whereas homozygotes (*db/db*) are severely obese and infertile, heterozygote females retain fertility and have 2.4-fold higher fat mass compared to age-matched wild-type control females (5). Thus, this model produces a consistent and reproducible maternal metabolic phenotype that can be used to examine effects of the intrauterine environment by studying wild-type offspring. Previously, we showed that Het_*db*_ pregnancy accelerates adipose accumulation and adipocyte maturation in early life and predisposes to diet-induced adipose tissue dysfunction (5). The development of adiposopathy or adipose tissue dysfunction is thought to be a central pathogenic event in obesity-associated metabolic disease. Healthy adipose tissue remodeling, driven by hypertrophy of existing adipocytes and generation of new adipocytes from a resident pool of adipocyte progenitors, ensures adequate buffering of excess energy under obesogenic conditions. A failure in adipose expansion to meet increased demands for lipid storage results in adipocytes that are excessively hypertrophic, inflammation, and impaired insulin-mediated inhibition of lipolysis, leading to lipid spillover into the circulation (7). Therefore, adiposopathy is a key mechanism linking obesity to NAFLD.

Results of the current study show that both male and female offspring born to Het_*db*_ dams are more susceptible to diet-induced obesity. Fasting levels of circulating FFA were higher in male offspring from Het_*db*_ dams when on a low-fat diet, with no further increase after prolonged high fat/fructose feeding despite a greater gain of whole-body fat mass. In contrast, female offspring from Het_*db*_ and normal pregnancies were protected from diet-induced increases in fasting FFA despite exaggerated diet-induced fat gain in Het_*db*_ female offspring. These sex differences were also observed in our previous study, which showed that female offspring were protected from increases in post-prandial levels of FFA after exposure to high fat diet (5). These findings led us to anticipate a sex-specific relationship between maternal adiposity and hepatic steatosis in the offspring. Contrary to our expectations, females and males born to overweight dams were similarly susceptible to diet-induced hepatic steatosis, despite maintenance of normal circulating FFA levels in females. Thus, the effect of an abnormal intrauterine metabolic milieu on risk for diet-induced hepatic steatosis in females appears to be uncoupled from circulating FFA levels, which are strongly correlated to DNL and fat accumulation in the liver (8–10).

Given the lack of sex differences in diet-induced hepatic steatosis, we undertook a lipidomics approach to investigate lipid metabolism in younger male offspring born to overweight dams. Profiles of FA species in FFA and TG extracted from the liver revealed diet-induced changes in the lipidome consistent with NAFLD-associated changes reported in the human and animal literature (10). Diet-induced increases the rate of DNL, as reflected by an increased ratio of palmitic acid to linoleic acid, were accompanied by increases in the abundance of SFA (palmitic acid and stearic acid) and MUFA (palmitoleic acid and oleic acid). Diet-induced increases in stearic acid were amplified in offspring from Het_*db*_ pregnancies, possibly due to increases in the elongation of palmitic acid observed in this group. The desaturation index (16:1/16:0), reflecting the conversion of SFA to MUFA for incorporation into complex lipids, was higher in high fat fed offspring from normal pregnancies. In contrast, HFFD decreased the desaturation of 18:0 to 18:1 in both groups of offspring, which was higher in Het_*db*_ offspring on a CD. While both ratios are thought to reflect SCD1 activity, the 16:1/16:0 ratio is more closely associated with DNL and NAFLD. A lower abundance of PUFA is the most common finding in NAFLD (10,11) and was also observed in high fat/fructose-fed offspring. The intrauterine environment had the greatest impact on PUFA levels, which were decreased in Het_*db*_ offspring fed a low fat diet or a high fat diet. PUFA levels are protective against hepatic steatosis and lipotoxicity as they have an inhibitory effect on lipogenesis and inflammation (10–12).

Collectively, these results support existing human literature demonstrating high pre-pregnancy BMI as a risk factor for development of NAFLD in the offspring. While an obesogenic phenotype was programmed in both male and female offspring born to overweight dams, diet-induced obesity in females occurred without changes in circulating FFA levels. Increased fat gain in response to high fat diet was associated with increased risk for hepatic steatosis in both sexes; however, male offspring displayed a stereotypic pathogenesis of NAFLD, arising from increased circulating FFA, whereas female offspring developed an equivalent degree of hepatic steatosis in the absence of changes to circulating FFA.

## Authorship Contribution Statement

J. A.T and T.B.S conceived and designed research questions; T.B.S., J.L.W., L.G.B., A.L.B., K. M.M., S.L., and J.A.T carried out experiments. S.U. performed histopathological scoring. T.B.S and J.A.T interpreted experimental results. T.B.S and J.A.T prepared figures. T.B.S and J.A.T drafted and edited manuscript.

## Acknowledgements

The authors would like to gratefully acknowledge the efforts of the staff at the University of Calgary Faculty of Vetrinary Medicine Diagnostic Services Unit for preparing samples for histology. We would also like to thank Dr. Jane Shearer for her invaluable input on the direction of this project. Finally, we would like to thank Dr. Stefan Urbanski for providing expertise in histopathological analyses.

